# Farm animal evolution demonstrates hidden molecular basis of human traits

**DOI:** 10.64898/2026.02.02.703413

**Authors:** Noah J. Connally, Shamil Sunyaev

## Abstract

Most human variants identified by genome-wide association studies are believed to affect traits by altering gene expression. This belief is supported by considerable circumstantial evidence, but statistical methods are unable to link most trait-associated variants to gene expression—a problem we refer to as “missing regulation.” Many explanations have been proposed, including the possibility that natural selection on gene expression limits power. Here, we take a novel approach to the question of missing regulation, beginning with the observation that the majority of trait-associated variants alter gene expression in two non-human species: cattle and pigs. We explain this discrepancy by comparing the species’ evolutionary histories. The observed differences in regulatory variants are consistent with selection on human gene regulation and increased genetic drift due to agricultural breeding. The differences are not limited to specific genes and reflect increased ascertainment of regulatory variants that are distal to genes. Additionally, we show that trait-associated gene regulation in cattle and pigs matches observed patterns from complex-trait genetics in humans, and may reflect currently unobserved trait-associated regulation in humans.

## Introduction

Although genome-wide association studies (GWAS) have identified tens of thousands of variant-trait associations in humans, the majority of these associations have not been linked to a molecular mechanism or gene. Most trait-associated variants are non-coding, and are often hypothesized to act by altering gene expression. This hypothesis is based on the enrichment of GWAS associations in regions of open chromatin, in regulatory elements such as enhancers, and among variants that alter gene expression (expression quantitative trait loci, or eQTLs)^1–9^.

The belief that GWAS would be explained by gene regulation helped motivate the creation of large, multi-tissue eQTL mapping datasets, along with numerous statistical methods for linking GWAS loci to eQTLs. Despite expectations, most GWAS variants cannot be tied to eQTLs^9–15^.

The disconnect between GWAS and eQTLs occurs across many different methods. Mediation methods estimate the fraction of a trait’s heritability explained by identified eQTLs; the fraction is less than 10-20%^16,17^. Transcriptome-Wide Association Studies (TWAS) measure the genetic correlation between GWAS and gene expression^18,19^. Roughly 36% of GWAS loci are correlated with the genetically predicted expression of gene^20^ (23% when corrected for confounders^21^). Colocalization methods test whether the causative variants for GWAS traits are also causally linked to gene expression (as opposed to two different variants that are in linkage disequilibrium). But colocalization explains only 8% to 42% of GWAS associations (see supplementary note 1 for the discussion of this wide range)^10,11,13–15,22–26^.

The limited ability to explain GWAS with gene expression persists even with known trait-relevant genes^13^, trait-relevant genes with eQTLs in relevant tissue^14^, and established drug targets^15^. For colocalization in particular, the success rate is roughly one-third what would be expected based on the enrichment of eQTLs in GWAS hits^25^.

We have used the term “missing regulation” to describe the seemingly small contribution of gene expression to GWAS traits. Due to the anticipated benefits of connecting GWAS and gene expression, there have been substantial efforts to explain why we do not find trait-associated eQTLs^14,15,12,27^.

Previous work suggests that natural selection may contribute to the problem of missing regulation. Analyses of GWAS show that selection shapes the genetic architecture of complex traits and affects which molecular mechanisms contribute to them^28–35^. A key signature of selection is an inverse relationship between variant effect sizes and frequencies, which occurs because many forms of selection prevent large-effect variants from reaching high frequencies. Evidence suggests that gene expression is also under selection^27,36,37^. However, selection can be difficult to test for molecular traits, whose effect sizes can vary across tissues, cells, and other contexts.

In fact, selection on gene expression might increase the context specificity of trait-associated eQTLs context specificity^12,27^. Methods for mapping eQTLs in specific cell types, cell states, developmental stages, or environmental conditions often lead to additional colocalizations with GWAS^38–49^.

Here, we set out to answer the following questions about the relationship between GWAS and gene expression.

First, are there numerous, undetected trait-associated eQTLs? Though natural selection and context-specificity may explain how trait-associated eQTLs could remain undetected, this does not demonstrate that they exist. The limited success in connecting GWAS to gene expression leaves open the possibility that other mechanisms are ultimately responsible^50^.

Second, what effects does natural selection have on the genetic architecture of gene expression? If selection were altered or absent, would eQTLs explain GWAS associations?

Third, if trait-associated eQTLs are widespread, how do they differ from ascertained eQTLs? Do they fall in different positions, different annotations, or differently selected loci?

When studying the effects of selection on traits, comparative genetics can offer insight. Recently, the Farm GTEx project has released eQTL mapping data for several agricultural species^51–54^. This has revealed an unexpected result: both cattle and pigs do not have the problem of missing regulation. In both species, GWAS results largely correspond to gene expression^51,52,55^.

In cattle and pigs, over 50% of GWAS heritability is mediated by eQTLs^52,56^ (vs. 15% in humans). TWAS connects over 90% of cattle and pig GWAS loci to gene expression^51,52^ (vs. 36% in humans)^20^. Using a consistent threshold for colocalization analysis, over 85% of GWAS associations in cattle and 55% in pigs are explained by eQTLs^51,52^ (vs. 42% in humans)^13,24^, all despite GWAS and eQTL mapping sample sizes (Figure 1).

**Figure 1.**
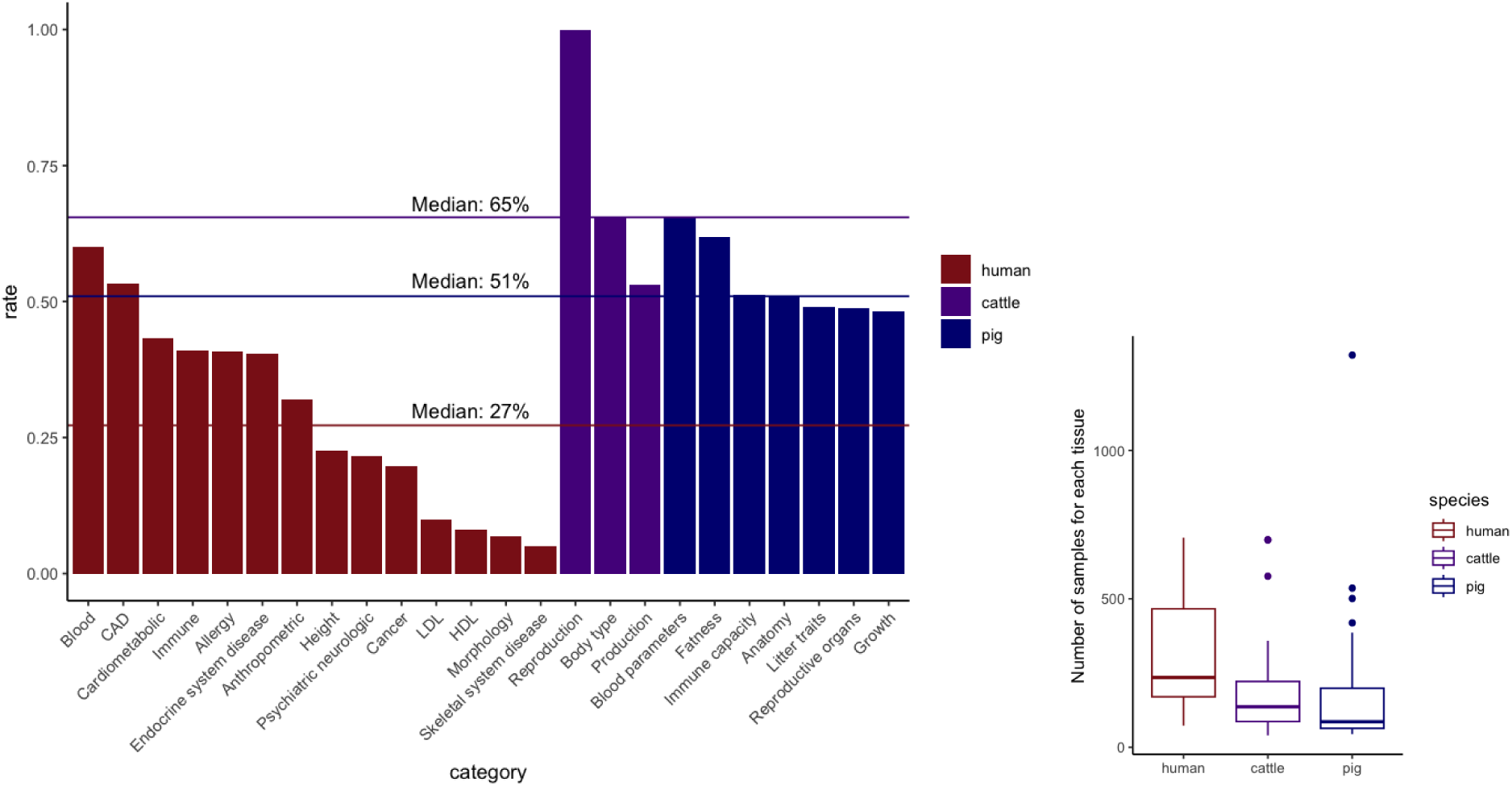
eQTL colocalization explains more GWAS loci in cattle and pigs than in humans. A) The fraction of GWAS loci explained using fastEnloc in three species: analysis of the human GTEx project v8^13,26^, the cattle GTEx project^51^, and the pig GTEx project^52^. Using a consistent posterior probability cutoff of 0.5, colocalization performs notably better in cattle and pigs. The horizontal bars represent the median across trait categories. Within humans, analysis of CAD, height, LDL, and HDL are from Hukku et al.; all others are from Barbeira et al. Categories were excluded if they had fewer than 20 GWAS loci to test. B) The number of samples across tissues in each of the three GTEx projects. Overall, human tissues have larger sample sizes, with some exceptions—most notably muscle tissue in pigs.

Humans, cattle, and pigs are all mammals, and we are not aware of evidence that they differ in their fundamental molecular biology. On the contrary, across these species similar traits are often associated with overlapping genes^57,58^ or even variants^59^.

One conspicuous difference between these species is their evolutionary history. Agricultural breeding in livestock has created strong artificial selection on a small number of traits. This strong positive selection has affected the entire genome, because even loci not under selection are affected by the genetic bottlenecks selection creates. These selection-induced bottlenecks, along with the bottleneck occurring in domestication, have altered the genetic architecture of cattle and pigs. It is possible that these differences explain why gene regulation explains GWAS so effectively.

In the paper, we compare the contribution of eQTLs to GWAS in humans, cattle, and pigs. We focus on the colocalization method fastEnloc^23,25^, which has been shown to have a low false-positive rate and an ability to work in the presence of allelic heterogeneity. We show that GWAS results are similar across species, but that human eQTLs are an outlier. Using simulations, we find that the discrepancy in colocalization success can be explained by the effects of evolutionary history on eQTL mapping. We argue that eQTLs observed in cattle and pigs indicate the existence of similar eQTLs in humans, and that those eQTLs make up much of the missing regulation.

### Evolutionary history alters colocalization success

Humans, cattle, and pigs are all mammals. Genes are strongly conserved across the three species, all have similar tissues, and the majority of agricultural traits studied are polygenic and driven mostly by non-coding variants^58^. However, the three species differ in their evolutionary histories.

Domestication and selective breeding of agricultural species have led to repeated genetic bottlenecks: timepoints at which only a small fraction of individuals contribute to the following generations. Genetic bottlenecks can result from a reduction in the total number of individuals (referred to as the “census population size”), but can also occur when genetic diversity decreases without a reduction in the number of individuals. “Effective population size” (N_e_) is a measurement of the random sampling of alleles that reflects genetic diversity—regardless of changes in census population size.

Though humans, cattle, and pigs all have large populations, their effective population sizes differ. During the last 150,000 years, the human N_e_ has increased five-fold or more^60–62^. Over the last 200,000 years, due to natural events, domestication, and artificial selection, agricultural species have experienced repeated bottlenecks. The N_e_ of cattle has fallen over 500-fold (for Holsteins)^63^, and the N_e_ of pigs has fallen over 300-fold (for Landrace, Yorkshire, Duroc, Hampshire)^64–66^. Estimates of current N_e_ (using common variation) are on the order of 10,000 for humans and 100 for Holstein cattle and major pig breeds^60–67^.

Critically for this study, a genetic bottleneck greatly increases the frequency of some deleterious alleles (while simultaneously removing many others). Trait-associated deleterious alleles at higher frequencies are more likely to be identified. Some studies improve statistical power by focusing on populations with bottlenecks. For example, some trait-associated variants have been identified only in Finnish ancestry, because the recent bottleneck of the Finnish population allowed these variants to reach higher frequencies^68–71^. However, the domestication and artificial selection of cattle and pigs has produced bottlenecks much stronger than any major human population. Because trait-associated variants and eQTLs are under selection, it is possible that similar variants exist at a much higher frequency in cattle and pigs than they do in humans^63,72,73^. If the difference in variant frequencies is substantial, it might explain differences in colocalization power.

We tested this by simulating two populations. Both populations have identical mutation rates, recombination rates, and mutation fitness effects. The values used are based on estimates from human genetics. The sole difference between populations was that one is based on the evolutionary history of western Europeans^61,67^ (the ancestry over-represented in GTEx) and the other on the evolutionary history of Holstein cattle^63^ (the most common breed in Cattle GTEx) (Figure 2A). (We do not separately simulate a population with a pig-based evolutionary history, given the overall similarity to the cattle evolutionary history.)

**Figure 2.**
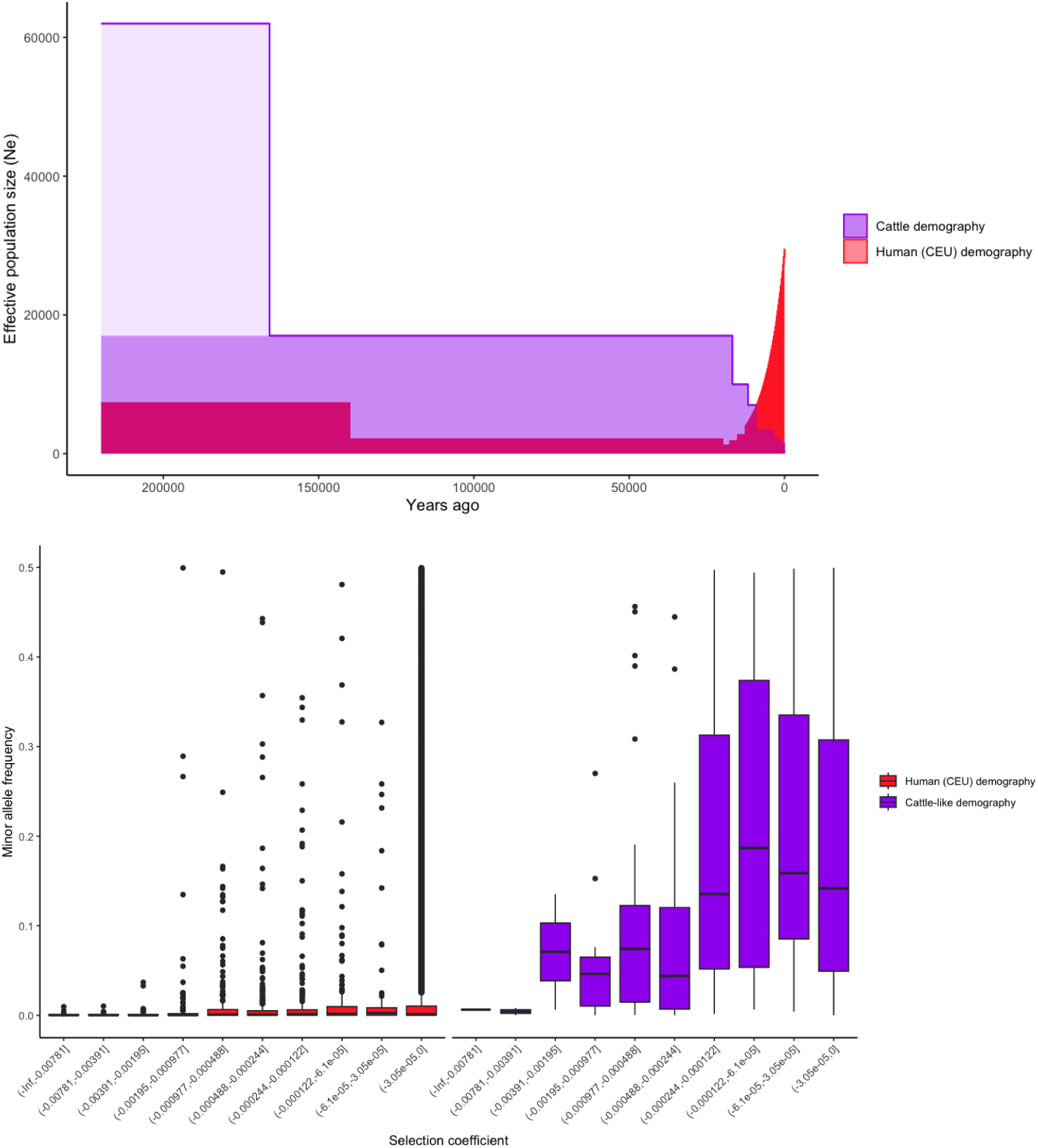
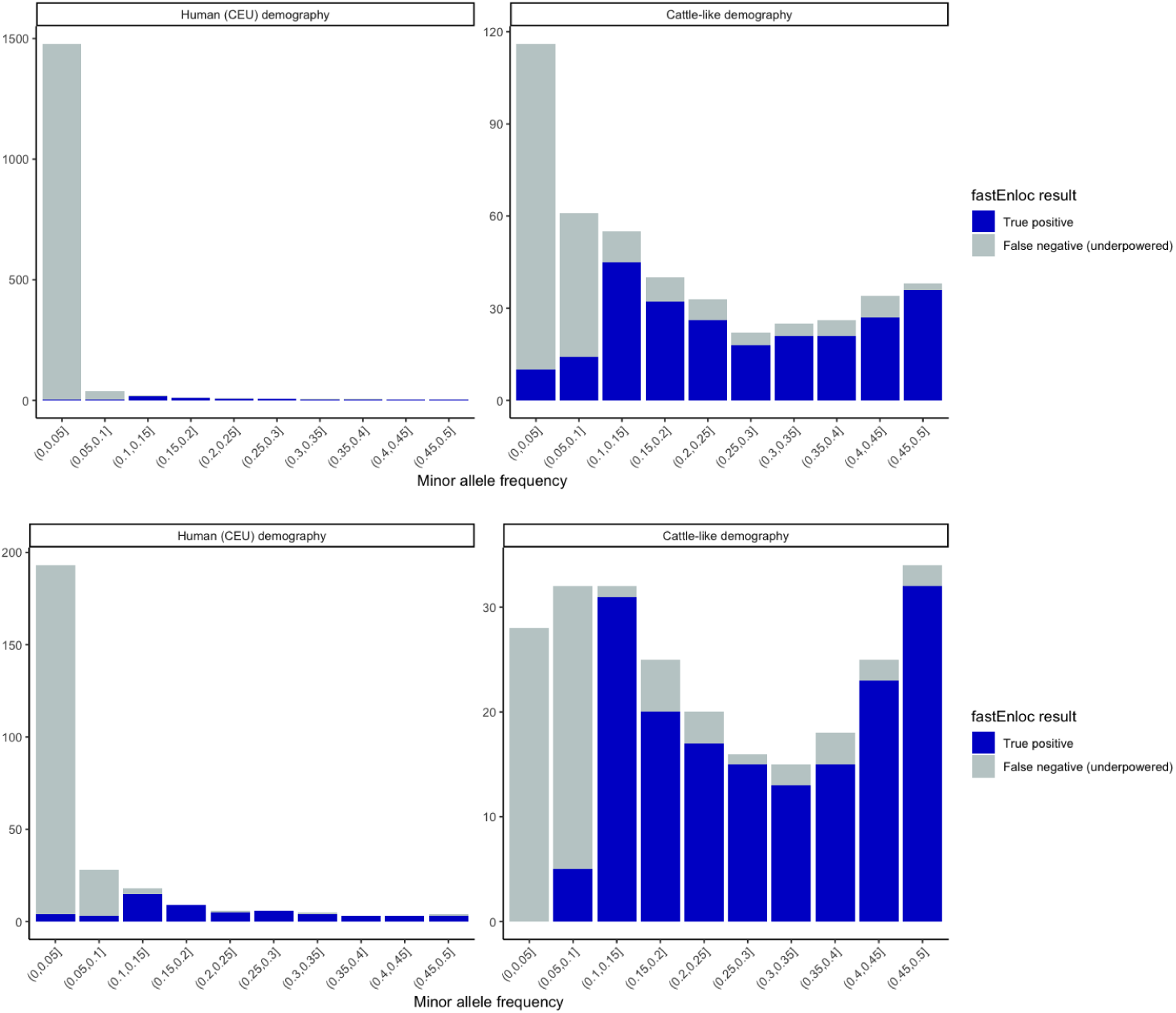
Changes in effective population size change colocalization power. A) The effective population size of humans and cattle over the past 200,000 years. The more pale shading within the cattle demography represents a portion of the cattle population history that we were unable to simulate due to computational constraints. B) The minor allele frequencies of simulated trait-associated variants. Each bar represents a different bin of selection coefficient, with the most deleterious variants on the left and the least deleterious on the right. Changing only the demography of the population produces large differences in allele frequencies. C) Colocalization success of different variants. Variants are grouped by minor allele frequency. The full bar shows the number of causative loci within each bin, while the blue portion shows the fraction of GWAS loci that successfully colocalize with gene expression driven by the same causative variant. D) As in C, but including only loci whose trait association could be detected in GWAS.

Following simulation, we examined the frequencies of variants under selection. The distribution of frequencies differed dramatically based on evolutionary history. Variants associated with complex traits have average selection coefficients of roughly −0.001 in humans^30,33^. In the population with a human evolutionary history, variants with approximately this selection coefficient had a median minor allele frequency (MAF) of 0.1%, while in the population with cattle evolutionary history, the median MAF was 14% (Figure 2B). This was consistent with the allelic ages of the simulated variants (Supplementary figure 2).

To simulate colocalization, we separated each population into GWAS and GTEx cohorts. Traits for both cohorts were simulated by assigning causative variants, and the effect sizes of variants were scaled based on their selection coefficients. We used fastEnloc to colocalize traits from the GWAS and GTEx populations. Because we know which variants are causative in the simulation, we can compare fastEnloc’s power in the two populations. In colocalization studies, the scaling of effect sizes and selection coefficients will vary, so this simulation is not meant to represent the absolute power of fastEnloc; its significance is the relative power in the two populations with different evolutionary histories.

Colocalization power differed dramatically between the two simulated populations. In the population with a European-based population history, 4% of causative GWAS loci correctly colocalized. In the population based on Holstein cattle population history, 56% of causative GWAS loci correctly colocalized (Figure 2C). The difference in power was driven by minor allele frequency of the eQTLs. These fractions represent the portion of all causative loci colocalizing. However, some loci are too rare, or their effects too small, to be detected by GWAS. When testing the portion of detectable GWAS loci colocalizing, the rates increase to 20% for the European population history and 70% for the Holstein population history (Figure 2D).

These simulations demonstrate that differences in evolutionary history can lead to large differences in colocalization success. When N_e_ is small and decreasing, genetic drift can bring trait-associated variants to high frequencies. This leads to increased colocalization power even without any changes in mutation rate, recombination rate, distribution of mutational fitness, or selection strength.

### Human bulk tissue eQTLs are under reduced selection

When natural selection acts on a trait, variants associated with that trait are selected on. As a result, finding that a variant is under selection increases the likelihood that it is associated with a trait. PhyloP is a method for measuring selection at each site in the genome, and variants at sites with higher PhyloP scores are more likely to be trait-associated in GWAS^74,75^.

PhyloP shows that human GTEx eQTLs are under weaker-than-average selection. This is inconsistent with human GWAS variants, and suggests that ascertained bulk-tissue eQTLs are unlikely to be linked with selected traits. However, in both cattle and pigs, bulk-tissue eQTLs are under stronger-than-average selection, consistent with those species’ GWAS and with a larger role for gene regulation in explaining complex traits (Figure 3).

**Figure 3.**
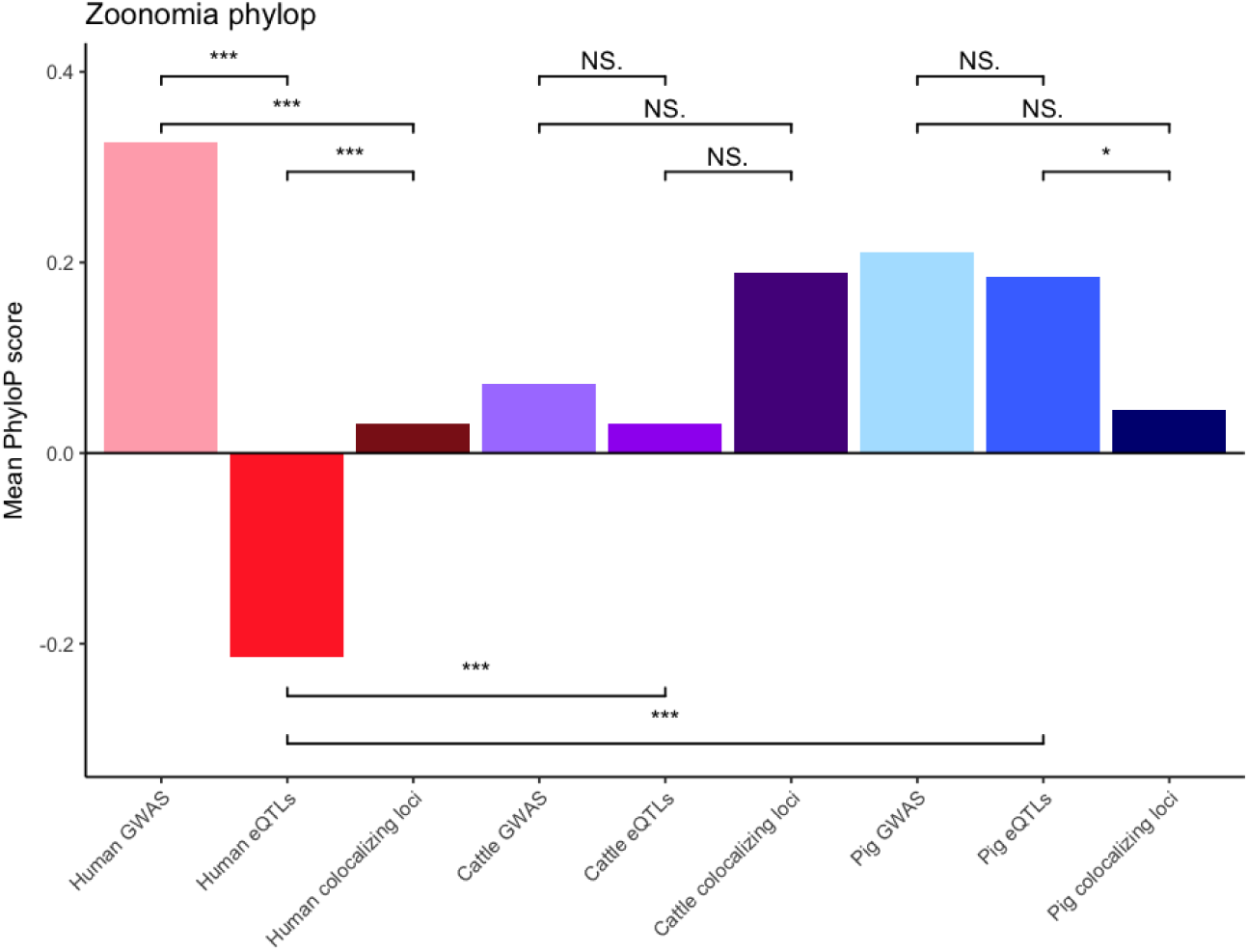
Human eQTLs have low evolutionary conservation. Each bar shows the mean evolutionary conservation of a set of SNPs, using the mammalian PhyloP scores from the Zoonomia Project^74,75^. A positive score indicates that the site is evolutionarily conserved—it is changing more slowly than would be expected by chance—which is often a marker of functional importance. Human eQTLs have, on average, negative PhyloP scores, indicating that they are less conserved than the average site in the human genome. In contrast, both cattle and pig eQTLs have positive scores, consistent with conservation. In all three species there is no evidence that GWAS variants or colocalizing loci have negative scores; human eQTLs are a statistically significant outlier when comparing against other human variants or against other species eQTLs. (“NS”, “*”, “**”, and “***” correspond to non-significant, P < 0.05, P < 0.01, and P < 0.001, after Bonferroni correcting for the 11 tests performed.)

### Cattle and pig eQTLs are consistent with “missing regulation” in humans

Human eQTLs are significantly enriched near the transcription start sites (TSS) of genes. GWAS variants have a much weaker enrichment near TSS; this discrepancy contributes to the low rates of GWAS-eQTL colocalization^27,76^. When colocalization does occur in humans, its TSS-proximity enrichment resembles eQTLs, not GWAS variants.

However, both cattle and pigs have smaller discrepancies between the TSS-proximity enrichment of GWAS variants and eQTLs. The distribution of distance from a GWAS variant to the nearest TSS is similar across all three species. But the TSS-distances of cattle and pig eQTLs resemble GWAS variants rather than human eQTLs (Figure 4A). As these distances might suggest, human eQTLs are substantially more enriched in annotated promoters than human GWAS variants, but both cattle and pigs have similarly low levels of promoter enrichment across eQTLs, GWAS variants, and colocalizing loci (Supplementary figure 1).

**Figure 4.**
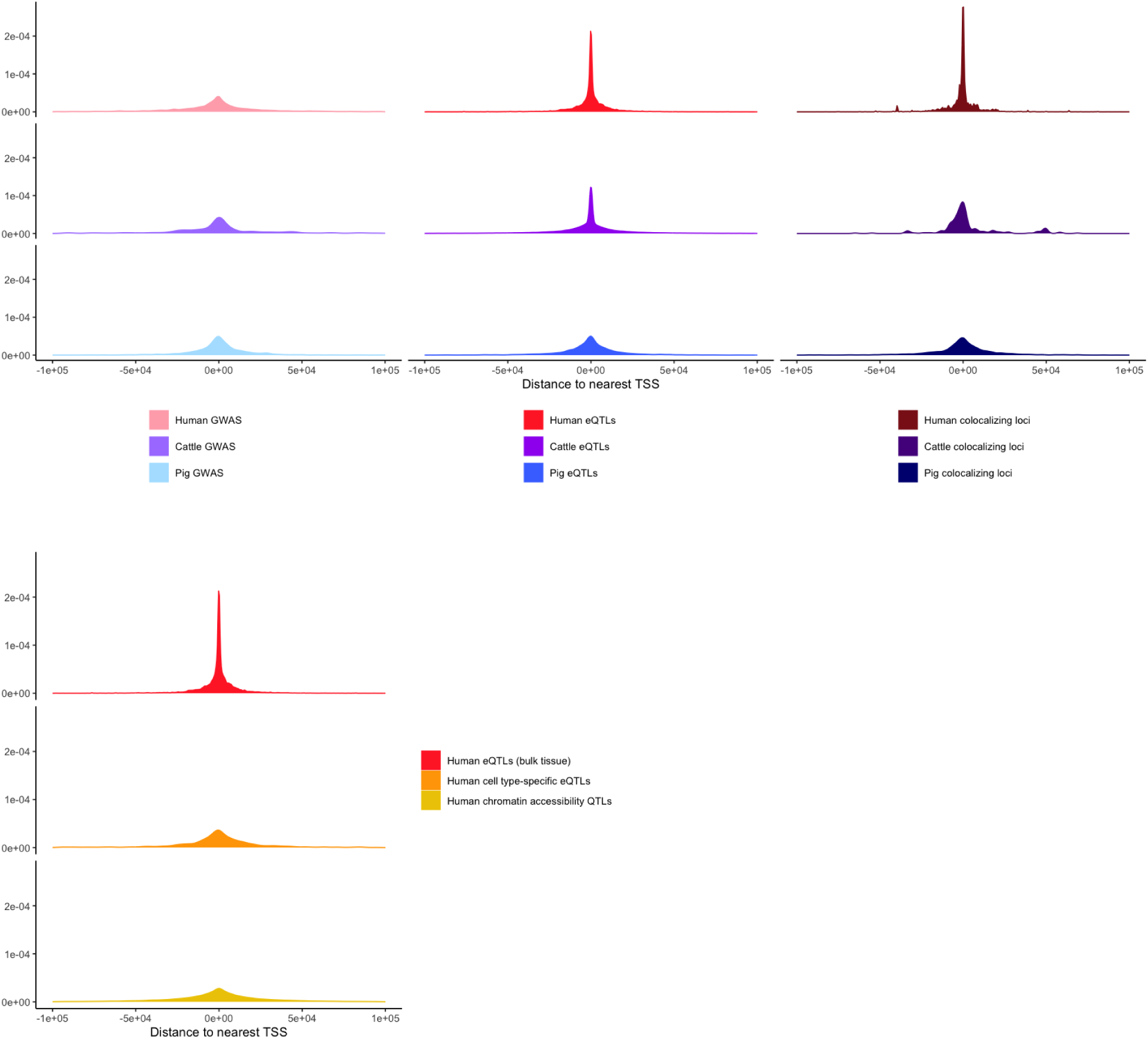
Human eQTLs are unusually close to transcription start sites of genes. A) Consistent with previous observations, we find that human bulk-tissue eQTLs are more strongly enriched near genes’ transcription start sites (TSS) than GWAS trait-associated variants are. Human colocalizing loci also have strong TSS-proximity enrichment. In cattle and pigs, GWAS variants have levels of TSS-proximity enrichment similar to GWAS in humans.

**Figure 5.**
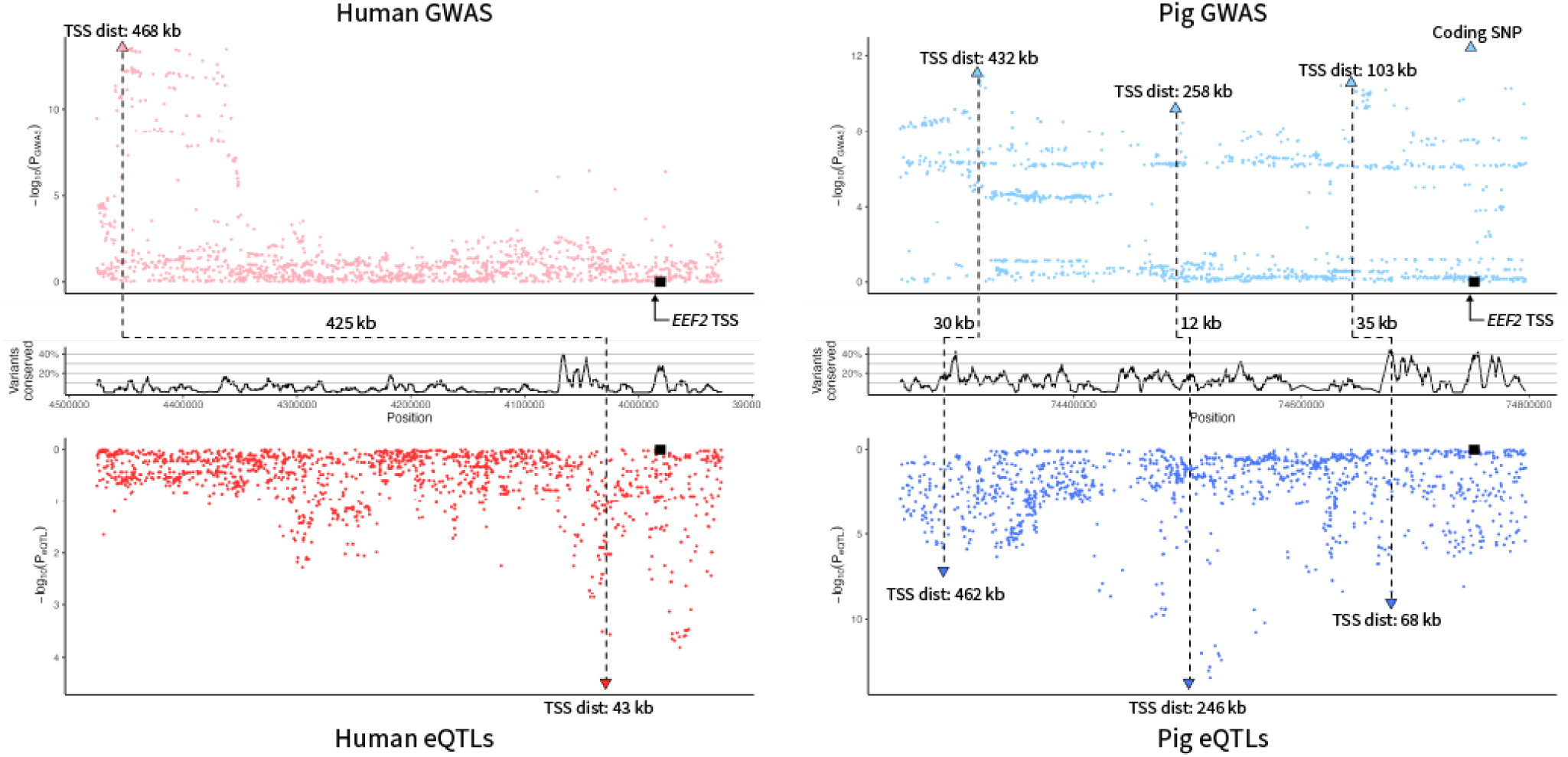
eQTLs explain GWAS loci near pig *EEF2*, but not human *EEF2*. In both humans and pigs, the gene *EEF2* has nearby GWAS hits for a trait related to musculature (fat-free body mass in humans, loin muscle depth in pigs). Both human and pig *EEF2* have eQTLs in muscle tissue. However, in humans, the GWAS lead SNP is over 400 kb away from the only eQTL. In pigs, there are three pairs of significant GWAS lead SNPs and eQTLs, each falling within 12-35 kb of one another. fastEnloc returns a negative result for the locus in humans, but a positive result in pigs for the most TSS-distal GWAS-eQTL pair shown above.

We also examined cell type-specific human eQTLs and human chromatin accessibility QTLs. With both of these molecular QTLs, GWAS traits co-localize at a higher rate than with bulk-tissue eQTLs. The TSS-distance distributions of cell type-specific and chromatin accessibility QTLs differ from human bulk-tissue eQTLs, but are similar to human GWAS They are also similar to bulk-tissue eQTLs in both cattle and pigs (Figure 4B).

However, in these species, bulk-tissue eQTLs and colocalizing loci have weaker TSS-proximity enrichment, and are more similar to GWAS. B) Unlike human bulk-tissue eQTLs, human cell type-specific eQTLs and chromatin accessibility QTLs both have lower TSS-proximity enrichment, similar to bulk-tissue eQTLs in cattle and pigs.

### Colocalization in cattle and pigs can identify genes missed by human colocalization

Cattle and pig GWAS are focused on agriculturally important traits. Many of these traits do not correspond exactly to any well-powered human GWAS, but some human and agricultural GWAS fall within similar categories, such as size or body composition. There are a limited number of traits that are measured in humans and an agricultural species. In available data, there are more corresponding traits in pigs than in cattle. For equivalent traits measured in humans and pigs, GWAS-eQTL colocalization in pigs can identify many additional genes potentially relevant to human biology. This is despite the smaller sample sizes of GWAS and eQTL studies in pigs.

The relative results of colocalization in humans and pigs depended on the traits being analyzed. For red blood cell count, substantially more genes had eQTLs colocalizing with the human GWAS than with the pig GWAS. This fits with the observation that blood traits exhibit high colocalization success using fastEnloc (Figure 1). For LDL cholesterol, more genes were identified through analysis in pigs than in humans. For traits related to size and body composition—important categories in animal agriculture—colocalization identified 1.7-fold and 3.9-fold more genes in pigs than in humans (Table 1; genes are listed in supplementary table 1). To test whether the genes identified in pigs were relevant to human traits, we examined the gene’s scores in PoPS, a method for prioritizing human genes based on data from GWAS, gene expression, protein-protein interactions, and biological pathways^77^. Genes identified in humans and pigs were similarly and significantly enriched in PoPS scores for relevant traits (Supplementary table 1).

**Table 1.**
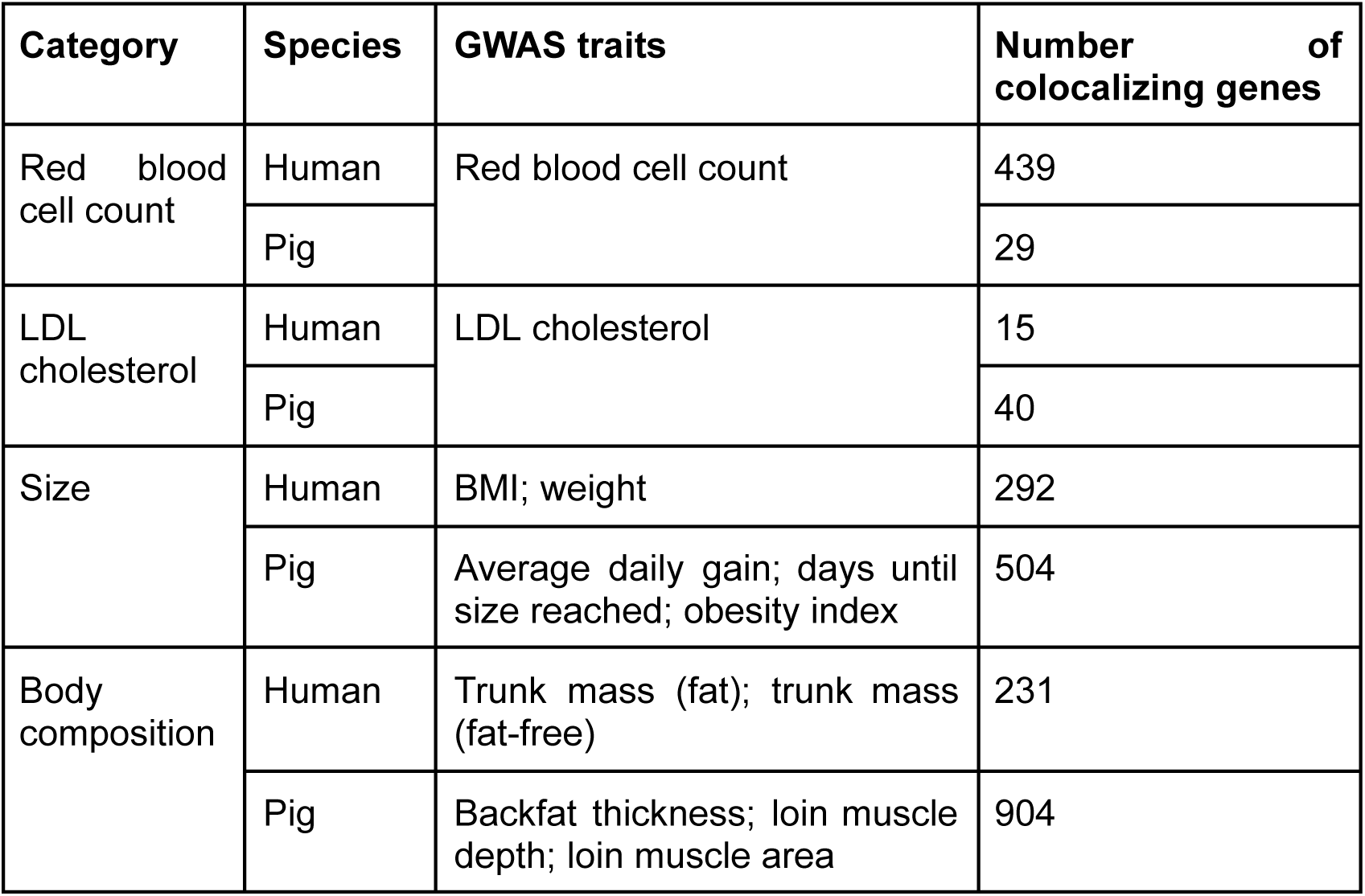
Number of colocalizing genes for corresponding traits in humans and pigs. We compare colocalization results for related traits in humans and pigs, looking at the number of genes identified in each species. Traits that directly correspond—red blood cell count and LDL cholesterol—we compare individually. Other traits we group by category. For example, body composition reflects the relative amounts of adipose and muscle tissue in an individual. In humans the trunk fat mass and trunk fat-free mass are relevant to body composition. In pigs, the relevant traits are backfat thickness and the depth and area of loin muscles.

Among genes identified with colocalization in pigs but not in humans, there are genes with well-established roles in biology of equivalent human traits. We have collected several notable examples. Burden testing links *INSR* (insulin receptor) to multiple anthropometric traits in humans, including fat-free weight, and distribution of adipose tissue^78^. Coding variants in *LRP5* (low density lipoprotein receptor-related protein 5) alter the distribution of adipose tissue across the body^79^, and burden testing links the gene to fat-free weight^78^. *PLIN4* (perilipin 4) coding variants can influence BMI, weight loss, adiposity, and the distribution of adipose tissue^80,81^, and are associated with weight and body composition through burden testing^78^. Each of these three genes—*INSR*, *LRP5*, and *PLIN4*—have mouse models with altered size and/or adiposity^82^. For each of these genes, fastEnloc colocalizes Pig GTEx eQTLs with body size and composition traits, but does not link their human GTEx eQTLs to similar traits.

*LDLR* (low density lipoprotein receptor) is a gene whose influence on LDL cholesterol levels is known through Mendelian disease^83^, animal models^82^, and burden testing^78^. In humans, fastEnloc found no colocalizations between *LDLR* eQTLs and the LDL cholesterol GWAS performed on the 361,141 samples in UK Biobank. LDL cholesterol did colocalize with an *LDLR* eQTL using fastEnloc v2.0^26^ and meta-analyzed data from the Global Lipids Genetics Consortium, a dataset with over 1.6 million samples^84^. In Pig GTEx, an *LDLR* eQTL colocalizes with a locus identified in an LDL GWAS of 860 pigs.

### Cross-species differences in gene regulatory architecture

To illustrate why fastEnloc is more successful in farm species, we compare *EEF2* (eukaryotic translation elongation factor 2) in humans and pigs.

*EEF2* was selected because it is an especially clear example of the effects we discuss (though the effects are general). It is strongly conserved, with few differences in the coding sequence between species. Previous research has shown that levels of active eEF2 protein influence skeletal muscle protein synthesis^85^, response to resistance training^86–88^, and muscle hypertrophy/atrophy^89,90^. Additionally, burden tests of rare coding variants have linked *EEF2* to multiple traits that are related to the size and proportion of musculature, including the mass and fat-free mass of the human body, torso, arms, and legs^78^.

Because eEF2 protein levels influence muscular development, it seems possible that *EEF2* eQTLs in muscle would colocalize with the mass or fat-free mass traits identified in GWAS. Muscle development is a trait of interest in pig agriculture, so we can also test for colocalization between *EEF2* eQTLs in pig muscle and traits such as loin muscle depth or area. In this instance, there is not a perfect match between traits measured in humans and farm species, so we must simply select GWAS in each species that reflect the same general trend: musculature.

Both species have genome-wide significant associations within 500 kb upstream of *EEF2*. In humans, the nearest fat-free mass association is 432kb upstream (Figure 4A). In pigs, there are several genome-wide significant loci, one falls within the gene, while the others are 68kb, 103kb, and 461kb upstream (Figure 4B). Humans have a significant eQTL 43kb upstream, and a distinct, but sub-significant eQTL 310kb upstream (Figure 4A). Pigs have significant eQTLs 68kb, 246kb, and 462kb upstream (Figure 4B).

*EEF2* exhibits the problem that we often see when trying to link gene expression to traits. A gene has an adjacent GWAS peak, but the peak is much more TSS-distal than the nearest eQTL (400kb more distal). But in pigs, we observe both a GWAS hit and an eQTL over 400kb upstream of the TSS. Though the GWAS and eQTL loci do not share a singular lead SNP, they colocalize, indicating that the same causative variant(s) underlies both.

This specific example is consistent with what we have observed broadly. Human lead variants for *EEF2* eQTLs are in regions with weaker selection than lead eQTLs in pigs. This may be because the evolutionary history of farm species has allowed high-frequency variation at these constrained sites. Both humans and pigs have a GWAS association nearly a half-megabase upstream. However, we find more TSS-distal *EEF2* eQTLs in pigs. As a result, we observe colocalization in the relevant tissue for pigs, but not humans

## Discussion

Linking GWAS and gene regulation can reveal trait-associated genes and the mechanisms behind non-coding GWAS hits. This potential is restricted by “missing regulation”—our limited ability to explain GWAS associations with eQTLs. There are multiple proposed explanations for this gap. Some explanations involve natural selection, which has been shown to influence the genetic architecture of complex traits.

Trait-associated eQTLs in cattle and pigs may help us understand the role of human gene expression in complex traits. But their relevance requires that the high success rate is not confounded by differences between species or methods that are irrelevant to our analysis. We have considered four possible confounders: linkage disequilibrium, imputation, positive selection on traits, and multi-breed GTEx.

The evolutionary history of cattle and pigs has altered linkage disequilibrium (LD) in the genome, which has the potential to influence colocalization. The decreasing effective population size leads to occasional high levels of long-range LD, but lower short-range LD, which is residual from the large ancestral population^91^. It is difficult to control for all sources and effects of LD. However, we have attempted to account for the effects of LD through simulation and assessing the trait-relevance of colocalizing genes.

Though we did not include them in our analysis, we have considered other potential confounders: imputation, positive selection on traits, and genetic diversity within eQTL mapping (Supplementary note 2).

In this work, we highlight evidence that eQTLs have the potential to explain complex traits, and show that trait-associated eQTL architecture in cattle and pigs differs substantially from the majority of ascertained human eQTLs. Our analysis supports the role of natural selection, while highlighting the importance of considering population history.

Non-human species have been used in the study of missing regulation because they expand the range of eQTLs that can be measured—for example, developmental eQTLs^92^, or eQTLs responding to manipulation of diet^93^. But to our knowledge, no studies have taken advantage of the possibility of comparing colocalization success across species in order to observe the effects of selection and evolutionary history. Colocalizations in other species can reveal information relevant to human biology, and different species may offer different benefits. For example, cattle and pigs have favorable variant frequencies, baboons have higher heterozygosity of eQTLs^94^, and our close relationship to chimpanzees enables us to search for human-specific gene regulation^95^. Even species without transcriptomic data can be useful. Cross-species differences in epigenomic marks correlate with differences in gene expression^96^, sequence conservation can be an indicator of functional regulatory effects^75,97,98^, and multi-species alignments can provide additional data for predicting regulatory changes^99^.

Though our results suggest that human GWAS may be explained by gene expression, they also indicate the eQTLs needed to explain traits may differ substantially from the majority of ascertained eQTLs in humans. Broadly speaking, there may be eQTLs that are trait-associated and eQTLs that are trait-independent. In this model, studies are well-powered to ascertain trait-independent eQTLs because selection tolerates high allele frequencies and activity over broad contexts, while trait-associated eQTLs are kept at low frequencies or act only in narrow contexts. As a result, most eQTLs ascertained in bulk human tissue are trait-independent, which explains their many differences with GWAS variants. Ascertained eQTLs are proximal to transcription start sites (TSS), while most GWAS variants are TSS-distal.

eQTL mapping in cattle and pigs has found eQTLs that are more TSS-distal, more strongly selected, and more likely to be trait-associated than eQTLs in humans. Because the population history of cattle and pigs is more favorable to mapping eQTLs, we suspect that these observations indicate the presence of similar eQTLs that exist in humans but are not widely detected in bulk tissue analysis.

If missing regulation is driven by context specificity, there may be further possible inferences about human eQTLs enabled by recent cattle and pig datasets that included addition types of data, such cell type-specific eQTLs, allele-specific expression, and cell type-abundance QTLs80,100,101.

This analysis supports the idea that natural selection contributes to missing regulation. In complex traits, selection limits the frequencies of variants with the largest effect sizes, while eQTL frequencies are limited based on their effect size, the impact of the affected gene, and the relevance of the contexts in which they act^12,27^. Our model could be improved by incorporating traits—including those under positive selection due to agricultural breeding.

However, our work shows that we should consider not only the selective pressures on a species, but also the species’ evolutionary history. Despite not modeling selection on specific traits, our simulations recapitulate both the absence of ascertained trait-associated eQTLs in humans, and their presence in cattle and pigs. The simulations showed that historical changes in effective population size could alter colocalization success independent of any changes in recombination rate, mutation rate, or sample size. The rapidly decreasing effective population sizes of cattle and pigs make them unusually well-suited for observing the effects of gene expression on traits. Humans, whose effective population size is increasing, may be an unusually poorly suited species.

We began this project with two observations. First, despite substantial evidence that gene expression influences complex traits, most human GWAS loci cannot be interpreted as eQTLs. Second, in both cattle and pigs, despite lower sample sizes, eQTLs colocalize with the majority of GWAS loci. By studying trait-associated eQTLs in other species, we can make inferences about human eQTLs. And by comparing these species—which have similar biology but different histories—we can examine the effects of evolution. We believe our results provide evidence for the existence of undiscovered trait-associated eQTLs in humans, and show the role of evolution in humans’ “missing regulation.” However, we have used only three species, and have barely scratched the surface of what a comparative approach to combining GWAS and gene regulation might uncover.

## Supplementary methods

### Comparison of results across species

We selected five studies performing colocalization using fastEnloc in humans, cattle, or pigs^1–5^. The cutoff for identifying colocalizations (regional colocalization probability, or RCP) differed between studies, so we compared all studies at RCP > 50%.

### Simulations of human and cattle demographies

We performed forward simulations using the program slim^6^ and the python library stdpopsim^7,8^. Parameters were selected based on Tennenssen *et al.* and Fu *et al.* for humans^9,10^, and on MacLeod *et al.* for cattle^11^. Because the effective population size of Holstein cattle is low, both population size and time were scaled up in order to sample more individuals. The resulting size prevented us from efficiently simulating the original cattle population, so the simulation began after the first bottleneck. The starting population at this size was simulated by creating neutral variation using msprime^12^, then changing the selection coefficients and running the simulation forward until the allele frequencies reached equilibrium.Neutral variation was added using python library tskit^13^ and msprime^12^. For both populations, we sampled 8,000 individuals for GWAS and 1,000 individuals for eQTL mapping. GWAS with 8,000 subjects are unusually small in humans and agricultural species, but computational constraints prevented us from using larger sizes. Phenotypes were simulated from the population using tskit^13^ and tstrait^14^. Plink 2^15,16^ was used to run GWAS.

### FastEnloc

We performed fine-mapping with dap-g^17,18^, using simulated summary statistics and individual-level genomes. We used fastEnloc^19,1,20,3^ to perform colocalization, with each colocalization including a 1 Mb region centered on the fine-mapped GWAS lead SNP. fastEnloc 2 can measure locus colocalization probabilities (LCP) rather than the regional colocalization probabilities (RCP). LCPs have higher power, but we used RCP for consistency with the previous analyses in the three species’ GTEx projects. We emphasize that simulations are only meant to compare the relative success between different demographies, not to infer absolute power under any scenario.

### Annotation enrichment

Promoters and enhancers were from application of ChromHMM to humans^21^, cattle^22^, and pigs^23^. In humans, the 18-state model was used in order to have higher similarity to the analyses in other species. Distance to transcription start site and enrichment in promoters and enhancers were measured using fine-mapped GWAS from humans^24^, cattle^25^, and pigs^5^, and fine-mapped eQTL data in humans, cattle, and pigs.

## Supplementary notes

**Supplementary note 1: Measuring colocalization success**

Published colocalization methods have used a variety of posterior probability cutoffs, ranging from 0.05% to 90%. This can contribute to large variation in the reported percentage of GWAS peaks colocalizing with eQTLs.

Additionally, in some GWAS-eQTL colocalization analyses, the genome is first divided into linkage-independent blocks. For the sake of example, let us consider height.

This approach often uses the blocks defined by. Berisa *et al.*^26^. Each block with a significant GWAS result is considered height-associated. Each height-associated block is tested for colocalization with eQTLs in each tissue. The success rate of colocalization is reported as the number of blocks with colocalizations divided by the number of height-associated blocks.

A weakness of this approach is that an individual block can have multiple independent associations to a trait. For a population of primarily European ancestry, there are 1,725 blocks across the genome, meaning the average block is roughly 1.7 Mb. For a highly polygenic trait such as height, it is almost guaranteed that some blocks will have independent associations (over 600 of the 1,725 blocks are height-associated). But each block is counted as a binary: colocalizing or not colocalizing. If a block has three independent height associations, and one of those associations colocalizes with an eQTL, the block is considered a colocalization success. If each block followed this pattern, it would be possible to report 100% colocalization success, despite only ⅓ of GWAS associations colocalizing with an eQTL.

A consequence of this method is that traits with more GWAS associations will have higher rates of colocalization, independent of any differences in their regulatory architecture. Traits with more GWAS associations also, by definition, contribute more of the tested loci. This can further inflate the success of colocalization. This effect is likely to be stronger in humans than in cattle or pigs, because humans have more GWAS with many loci. Though we are not certain, it may contribute to the difference between the total rate of colocalization (42%) and the median rate of colocalization per trait (21%) in the human GTEx v8 data (Supplementary figure 30 of ref^27^).

**Supplementary note 2: Consideration of possible confounders**

The evolutionary history of cattle and pigs has altered linkage disequilibrium (LD) in the genome, which has the potential to influence colocalization. The decreasing effective population size leads to occasional high levels of long-range LD, but lower short-range LD, which is residual from the large ancestral population^28^. It is difficult to control for all sources and effects of LD. However, we have attempted to account for the effects of LD through simulation and assessing the trait-relevance of colocalizing genes.

In the human GTEx project, RNA-sequencing and whole-genome DNA-sequencing were performed on donors^27^. In the cattle and pig GTEx projects, RNA sequencing was performed, and genomic regions adjacent to genes were imputed from the RNA sequencing. Imputation accuracy was high (when tested against a genotyping array), but not perfect, especially for genes with lower expression^5,29,30^. However, to produce false positives, variants would have to be mis-imputed in ways that systematically occurred at the same positions as GWAS hits and were correlated with inter-individual differences in gene expression. This seems unlikely, and we do not believe imputation errors contribute to the high colocalization success of cattle and pigs.

GWAS in cattle and pigs are focused on a limited number of commercially significant traits. Because of their importance, these traits are subject to strong directional selection, which can change allele frequencies and linkage disequilibrium^31–33^. However, we note that cattle and pig eQTLs differ in genetic architecture from human eQTLs across the entire genome, not just for genes related to agricultural traits. This points to differences that extend beyond the specific traits analysed.

The cattle and pig GTEx projects are multi-breed, while the human GTEx project primarily has subjects with European ancestry. Cohorts with greater genetic heterogeneity can improve GWAS or eQTL-mapping power, whether the heterogeneity comes from more diverse human ancestries^34–38^ or multiple agricultural breeds^39–41^. Genetic heterogeneity can improve power for colocalization and other methods of linking eQTLs and GWAS^42,43^. Our simulations show that the evolutionary history of species can explain large differences in colocalization, even without including multiple ancestries. Nonetheless, diversity of ancestry may be an additional contributor to colocalization power in cattle and pigs.

In our simulations, GWAS loci for the cattle demographic history were more likely to colocalize with multiple eQTLs than GWAS loci from the European demographic history were. This may be true for causative loci in real cattle and pigs, though our simulations contained an unrealistically high density of eQTLs. Regardless, even when loci with high numbers of colocalizations were removed, the power of fastEnloc remained higher with the simulated cattle demographic history.

## Supplementary figures

**Supplementary figure 1.**
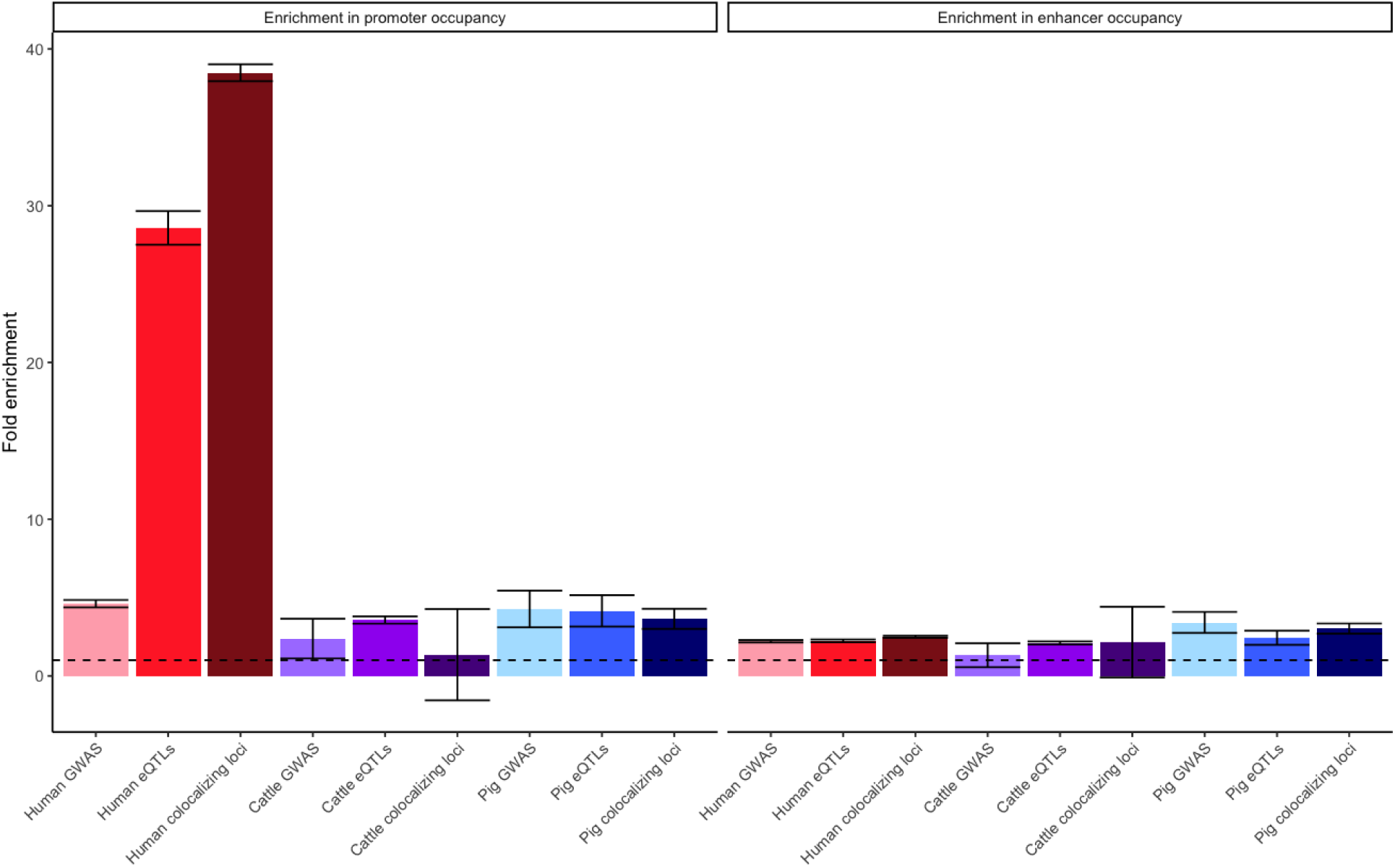
Human eQTLs are unusually promoter enriched. Human eQTLs have strong enrichment in promoter occupancy. This differs not only from human GWAS, but also from eQTLs in cattle and pigs. We did not observe clear differences in enhancer occupancy enrichment across species.

**Supplementary figure 2.**
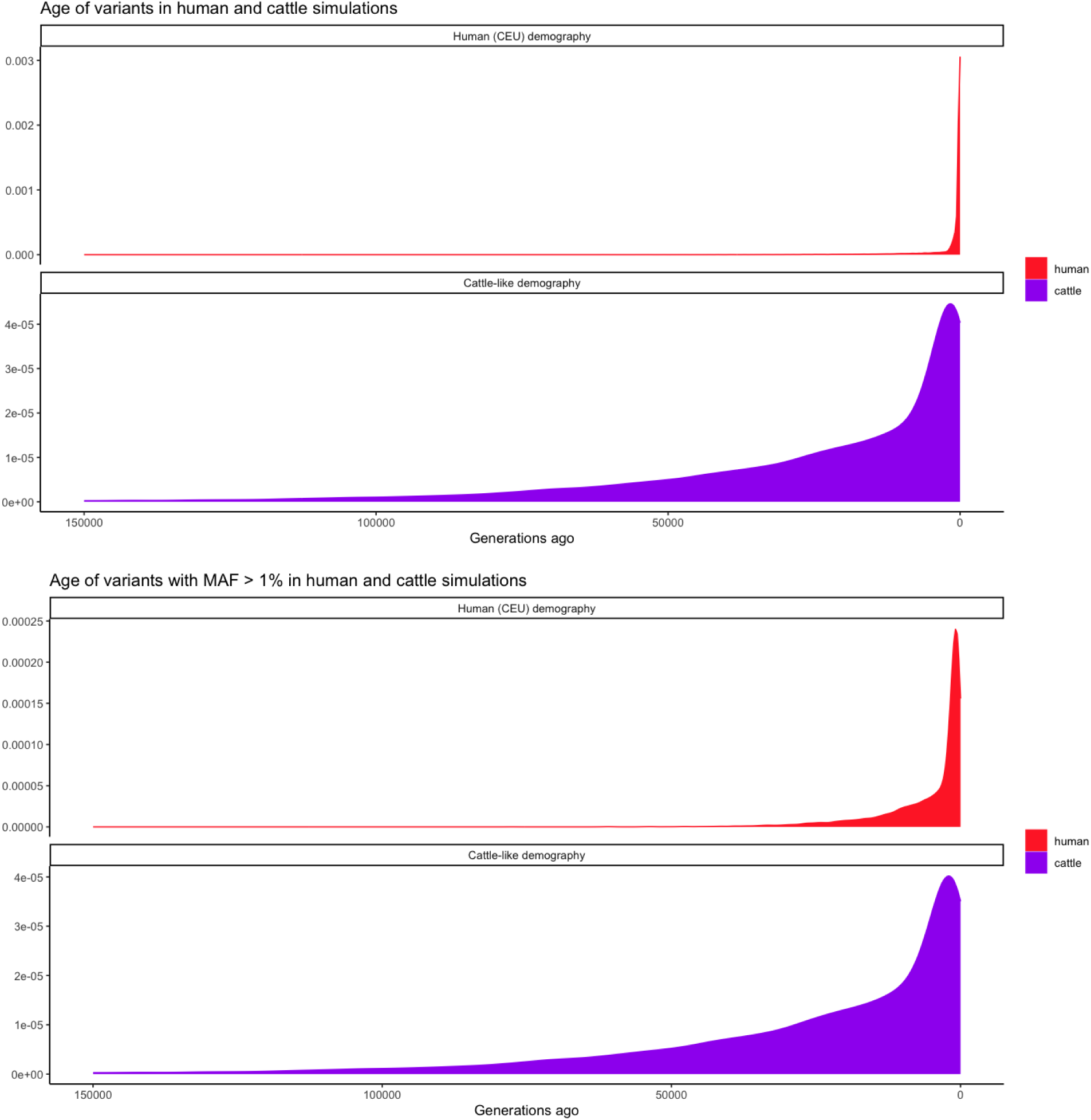
Allelic ages of simulated variants. We recorded the allelic ages of variants present in the final generation of our simulated populations. A) Allelic ages were larger in the population based on the cattle demography history. B) This age difference remains even when considering only variants with a minor allele frequency of at least 1%.

## Supplementary tables

**Supplementary table 1.**
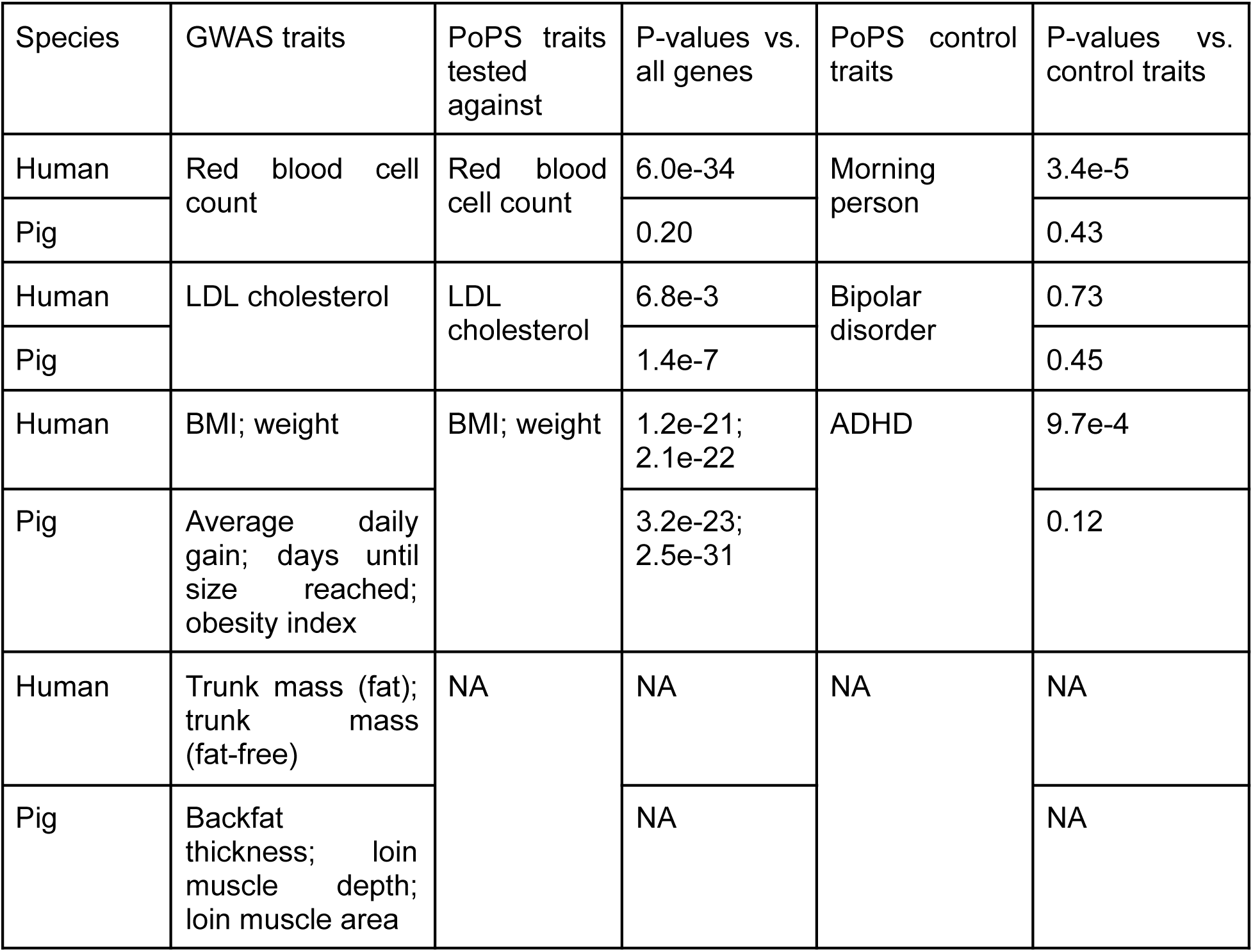
Trait-relevance of genes identified by fastEnloc. For each set of colocalizing genes, we tested for enrichment in PoPS scores^44^ for relevant traits. P-values were calculated with a T-test comparing the PoPS scores of colocalizing genes to all other genes. It is possible the genes relevant to one trait are enriched not because of a specific trait relationship, but because they are broadly important genes. To confirm that this does not drive our results, we ran an additional T-test comparing colocalizing genes’ PoPS scores in a relevant trait to their scores in a trait with no clear relationship (“control trait”). In every case in which colocalizing genes were enriched in PoPS scores, they remained enriched when comparing the PoPS scores of the relevant trait to the scores for the control trait.

**Supplementary table 2: Genes identified by colocalization in humans and pigs.**

This table is included as the “file supplementary_table_2.tsv” We report which genes colocalized for each of the GWAS traits included in table 1 of the main text. For human genes we use results from the studies conducted with the original version fastEnloc^1,2^, but not from Hukku *et al.*, as fastEnloc v2.0 has a “locus-level colocalization probability” is difficult to compare to the “region colocalization probability” used by the other studies^3^.

## Data and code availability

Code for conducting the analyses in this paper is available at https://github.com/NJC12/colocalization_humans_cattle_pigs. The study used only publicly available data, and links to download those data are included in the analyses.

## Acknowledgements

This work was supported by NIH grants R35GM127131, R01MH101244, and U01HG012009.

Many people have been generous with their advice for this project, and we would like to thank (in alphabetical order) Carles Boix, Colby Chiang, Ran Cui, Jared Decker, Steven Gazal, Dailu Guan, Elinor Karlsson, Evan Koch, Aoxing Liu, Aparna Nathan, Luke O’Connor, Stephen Riley.

